# Prediction of Polygenic Risks by Screening Thousands of Polygenic Scores

**DOI:** 10.1101/2025.10.03.680330

**Authors:** Wei-Min Chen, Ani Manichaikul, Suna Onengut-Gumuscu, Bradford B. Worrall, Stephen S. Rich

**Author notes:** Corresponding Author: Dr. Wei-Min Chen, P.O. Box 800717, Department of Genome Sciences, University of Virginia, Charlottesville, VA 22908 USA.

## Abstract

**Rationale:** A polygenic score (PGS) summarizes a person’s genetic information in a single number for a trait, and its utility in genomic medicine is well-recognized. Although PGS models have been generated for many traits, they are not broadly available for certain traits due to limited sample sizes in studies of infrequent outcomes. Often, prediction of these not well-studied traits (*e.g.*, treatment/drug responses) would be of great clinical utility and have been underutilized due to statistical power limitations. To support more versatile trait prediction, we present a method of developing PGS models that can be used in studies of any size with genome-wide SNP data.

**Method:** We first generate thousands of PGSs for each study participant in a given data set using their genome-wide SNP data and public resources. A PGS-wide scan involves evaluating the Area Under the Curve (AUC) of prediction for a binary trait (or the R-squared of association for a quantitative trait) at each PGS. We present two methods for the PGS model development, SECRET-Best, which selects the most predictive PGS from the PGS scan for prediction, and SECRET-WTSUM, which considers a combined score from multiple correlation-pruned PGSs. This algorithm is scalable and implemented in a user-friendly software tool, SECRET (Screen and Evaluate Catalogued Risk scores to Enhance Trait predictions).

**Results:** We applied SECRET to a binary outcome (type 1 diabetes [T1D]) and a dataset of 2,100 samples, each with 12 laboratory test-related continuous traits. For nine traits with existing PGSs available in the PGS catalog, eight of the traits were predicted correctly, with the same trait-related PGS identified as the top predictor. We showed that the existing PGS methods had rather limited power to predict trait values in a validation set when only 1,500 samples were used to develop a PGS model, while the SECRET methods were able to maintain the prediction power for under-powered GWAS studies even when the sample size of the study was in hundreds.

**Conclusion:** The SECRET methods and tool provide a valuable resource for studies with genomic data but of limited sample sizes. This approach enables systematic development and evaluation of PGS models.

## Introduction

A polygenic score (PGS) aggregates the effects of many genetic variants into a single number and is predictive of an individual’s disease risk. The utility of PGSs in genomics and medicine has been well-recognized in recent years, with applications including disease risk prediction^1–3^, PGS-based screening of patients for treatments or drugs^4,5^, testing gene-environment interactions or shared etiology between diseases^6^, and applications in disease diagnosis, prognosis and subtypes^7^.

A PGS is typically calculated by summing the number of trait-associated alleles in an individual, with each weighted by its effect size as determined from a genome-wide association scan (GWAS). Various methods and tools^8–17^ have been developed to construct a PGS model for a trait. For many widely-studied traits, it is often possible to directly use an existing PGS model without needing to develop a new one. While PGSs are somewhat widely-used for common phenotypes and disease traits, they remain unavailable for less frequently measured traits such as treatment or drug responses. In such cases, generating a GWAS-based PGS model may not be practical due to limited sample sizes for infrequent disease outcomes or in a clinical trial study. Alternative ways of developing PGS models are needed in addition to relying on existing PGS models.

Thousands of publications on PGSs motivated the creation of public PGS resources such as the PGS Catalog^18^, the ExPRSweb^19^, and the Cancer PRSweb^20^. The PGS Catalog is the largest resource, containing 5,161 PGSs for 659 traits in its latest version. With the annotated scoring information and the curated metadata, these catalogs provide researchers with excellent resources to help generate PGSs for their data. Tools such as PLINK^21^ and VIPRS^22^ can be used to calculate PGSs from a PGS model in the catalogs once the GWAS data are imputed. More efficiently, one can obtain all 1,000s of PGSs for each study participant by simply uploading GWAS dataset to the Michigan imputation server^23^ and requesting the PGS calculation.

To allow for more versatile PGS applications, we present SECRET (Screen and Evaluate Catalogued Risk scores to Enhance Trait predictions), a method of developing PGS models that can be used in studies of any size with genome-wide SNP data. The concept is to leverage 1,000s of existing PGS models (such as those found in the PGS Catalog^18^) to identify the best PGS model that is predictive of the trait of interest. Since a dataset consisting of 1,000s of PGSs for each study participant can be analyzed similarly as genomes, the resulting dataset is a PGS-wide score data. Similar to a genome-wide association scan, a PGS-wide association scan involves evaluating the prediction power for the trait of interest at each PGS in the PGS-wide score data. SECRET is one of the first tools to leverage public resources to develop PGS models for predicting traits of interest. SECRET is both informative and powerful, especially when the initiating study has a small or modest sample size. We present the SECRET approach and use real data examples to illustrate the utility of SECRET in PGS applications.

## Methods

### Methodological Framework: Screen and Evaluate Catalogued Risk scores to Enhance Trait prediction (SECRET)

A PGS-wide association scan begins with evaluating the prediction power at each PGS in a PGS-wide score data. By scanning all the PGSs, we can identify the best predicting PGS models. For quantitative traits, the prediction power is quantified as the R-squared of association between a PGS and a trait to be predicted in a linear regression framework. Suppose *S_im_* is the regression residual of the *m-*th PGS for the *i-*th individual after adjusting for top PCs of ancestry. Suppose *Y_i_* is the regression residual of a trait for the *i*th individual after adjusting for top PCs of ancestry and other covariates such as sex and age. At the *m*-th PGS, the regression model is:

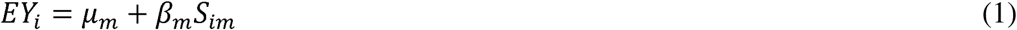

where *β_m_* is the regression coefficient. Let *S_m_* and Y be vectors consisting of *S_im_*’s and *Y_i_*’s respectively, with means 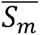 and *Y̅*, and variances 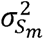 and 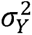 respectively. Assume the sample size is *N*. The test statistic for the linear regression analysis at the *m*-th PGS is:

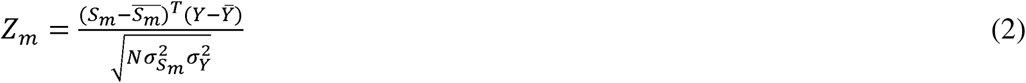

which follows a standard normal distribution under the null hypothesis of no association between the *m*-th PGS and the trait. The prediction power can be evaluated using r-squared between the *m*-th PGS and the trait:

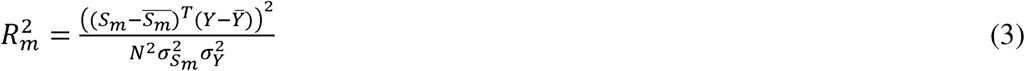

Note the 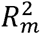 is the square of the correlation between the PGS and the trait. It is also the proportion of phenotypic variation that is explained by the PGS.

After the PGS-wide association scan, all PGS models can be ranked by their *r^2^* with the trait. One special PGS model for the trait is the PGS with the largest *r^2^*, and we name this PGS model S^Best^. Accordingly, we name this method SECRET-Best.

With the top-ranked PGS models identified from a PGS-wide scan, it is possible to further improve the PGS model by considering a composite PGS. To reduce duplicated information, we first prune correlations among the top-ranked PGSs and only consider PGSs that are not strongly correlated (*e.g.*, *r^2^* < 0.2) with those more predictive ones. One way to construct a composite PGS for the *i*-th individual is to consider the weighted sum of these top-ranked correlation-pruned PGSs, with the weight being the regression coefficient in the regression of the trait on the PGS:

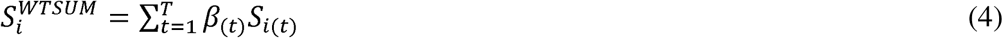

where *β*_(*t*)_ is the regression coefficient in Equation 1 at the *t*-th correlation-pruned PGS, *S_i_*_(*t*)_ is the *t*-th correlation-pruned PGS for the *i*-th individual, and *T* the total number of top-ranked correlation-pruned PGSs. We name this PGS method SECRET-WTSUM.

For binary traits in a case-control study, the PGS-wide scan involves evaluating the Area Under the Curve (AUC) at each PGS under a logistic regression model. Following the PGS-wide association scan, all PGS models are ranked by their AUCs with the trait. SECRET scores S^Best^ and S^WTSUM^ can be similarly developed.

### Extension of SECRET to Family Data

Family data provides unique opportunities for PGS applications, where confounding factors such as population stratification and assortative mating are well controlled. SECRET implements a family-based PGS method that can handle the parent-offspring trio data. Suppose in the *f*-th parent-offspring trio, the *m*-th PGSs for the father, the mother, and the offspring are 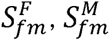, and 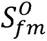, respectively. Then the adjusted PGS at the *m*-th PGS for the offspring is 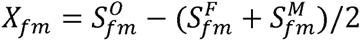. Let 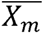 and 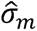 represent the average and the standard deviation of all adjusted PGSs at the *m*-th PGS, and *N* be the total number of trio families, then in affected offspring only we can carry out a TDT-like test:

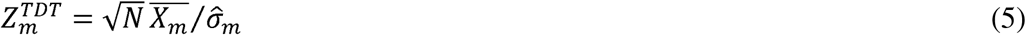

which follows a standard normal distribution under the null hypothesis of no association between the *m*-th PGS and the affection status. In the presence of unaffected offspring, we can also consider a case-control analysis:

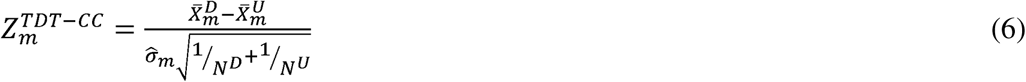

where *N^D^* and *N^U^* are the numbers of affected and unaffected offspring respectively, 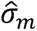 is the standard deviation of the pooled adjusted PGS, and 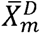 and 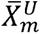 are the average adjusted PGS in affected and unaffected offspring respectively at the *m*-th PGS. With the adjusted PGSs calculated, the AUC to predict the case/control status at each PGS for the TDT-CC method can be computed in the same fashion as in the standard case-control analysis. The PGS with the largest AUC is the SECRET score S^Best^.

### Computational approach

We implemented all algorithms in a scalable and user-friendly software tool, SECRET. This tool takes PGS-wide score data and binary or quantitative traits as input, evaluates and identifies the most predictive PGS models for the trait of interest while adjusting for PCs of ancestry and other covariates. The PGS-wide score data required by SECRET can range from a single custom PGS to 1,000s of PGSs per person, as in the current PGS Catalog, or >10,000s, as in future versions of the PGS Catalog, providing users the flexibility to evaluate both catalog-based and custom PGSs.

SECRET is written mostly in C++, with some R code for visualization of prediction and for extra flexibility of method development (*e.g.*, the implementation of Cox proportional hazard regression model for time-to-event traits). Multi-processing using OpenMP and other efficient computation techniques^24^ allows computation of large-sized PGS-wide association scans in a matter of seconds.

### Data application: quantitative traits from the VISP study

To demonstrate the prediction power of the SECRET methods, we first used the Vitamin Intervention for Stroke Prevention (VISP) GWAS data^25,26^ to generate the PGS-wide score data using the Michigan imputation server^23^. The VISP trial was a clinical trial that investigated whether high doses of folic acid, vitamin B6, and vitamin B12 could reduce recurrent strokes in patients who had previously experienced a non-disabling ischemic stroke. The VISP data in this analysis consists of 2,100 samples of diverse ancestry (1,723 samples of European ancestry and 377 samples of non-European ancestry), each with 12 lab test-related traits measured, including 8 traits with existing PGS models available in the PGS Catalog (creatinine, total cholesterol, high density lipoprotein (HDL), triglycerides, vitamin B12, C-reactive protein (CRP), diastolic blood pressure (DBP), and systolic blood pressure (SBP)), and 4 traits without existing PGS available (factor XII (F12), vitamin B6, folate, and Von Willebrand factor (vWF)). The top 10 PCs of ancestry are adjusted for both PGS and the trait values in the linear model (Equation 1). Sex and age are additionally adjusted as covariates for the trait values.

We first carried out PGS-wide scans using SECRET in all 2,100 samples and then in 377 samples of non-European ancestry only. To evaluate the validity of prediction using the SECRET methods, and to compare the prediction performance of SECRET with existing PGS methods, we randomly split the 1,723 samples of European ancestry into a training set of size 1,500 and a validation set of size 223. With the training set, we developed 5 PGS models using 2 SECRET methods (SECRET-Best and SECRET-WTSUM), and 3 existing methods, including Clumping and Thresholding (C&T)^12,13^, Lassosum2^14,15^, and LDpred2^16,17^. All 3 existing PGS methods were implemented in the PennPRS webserver^27^. For the SECRET-WTSUM method, we applied a correlation filter *r^2^*<0.2 and used 10 top-ranked correlation-pruned PGSs. For PGS methods available through the PennPRS webserver, all parameters (such as the p-value threshold for the C&T method) were determined by a cross-validation procedure. We tested and compared the performance of the 5 PGS models in the validation set.

### Data application: Prediction of T1D in affected families

We applied the family-based PGS scan methods (Equations 5 and 6) to the T1D Genetic Consortium (T1DGC) family data^28^ consisting of 12,213 samples in 4,694 parent-offspring trios. Our family-based PGS-wide scan (Equation 6) considered offspring with both parents available (N=4,694), which included 3,734 offspring with T1D and 960 offspring without T1D.

Next, we explored pathways and etiology that T1D may share with other diseases, through the Major Histocompatibility Complex (MHC) region. We calculated the MHC PGSs using SNPs from the MHC region only and the non-MHC PGSs using SNPs outside of the MHC region. We ran an MHC and a non-MHC PGS-wide scan, separately.

### A simulation study to assess the sample size and power

To determine the sample size needed by SECRET, we conducted extensive computer simulations using the VISP and T1DGC datasets. In each round of simulation, we randomly selected a subset of samples (N=100, 200, 400, and 800 respectively) for 8 traits from VISP and T1D from T1DGC family data. For the family data, the random selection was in families, and we randomly selected N/4 families from 840 families consisting of two parents and two affected offspring only, *e.g.*, N=100 corresponds to 25 such families. We carried out a PGS-wide scan using SECRET and evaluated whether a PGS for the trait of interest could be identified as one of the top PGSs (*e.g*., top 1, top 3, or top 10). We repeated this procedure 1,000 times and calculated the power of identifying the correct PGS for the trait.

## Results

### Data application: matching traits to the most predictive PGS in VISP

Table 1 shows the most predictive PGS results identified from each of the 12 PGS-wide scans in VISP. For 8 traits with existing PGSs available in the PGS catalog, which included creatinine, total cholesterol, HDL, triglycerides, vitamin B6, CRP, DBP, and SBP, all traits but B12 were predicted correctly, with the same-trait-related PGSs being identified as the most predictive predictors. The best predictor for B12 is a PGS underlying the “Vitamin B12 deficiency induced Anemia” trait, which is closely related to the B12. The prediction *r^2^* between the best PGS and the trait ranged from 0.01 (for DBP) to 0.105 (for HDL). All 8 associations were PGS-wide significant, with the largest P=3.7 × 10^−6^ (for DBP). For the 4 traits without existing PGSs, the best predictors were closely related to the underlying biology of the trait. The Factor VIII levels PGS predicts vWF (Von Willebrand factor) with *r^2^*=0.071, the phosphate PGS predicts B6 with *r^2^*=0.017, the arthrosis of knee PGS predicts folate with *r^2^*=0.016, and the SBP PGS predicts factor XII with *r^2^*=0.008. All predictions were catalog-wide significant with P < 1.1 × 10^−5^ (Bonferroni threshold with 4,477 PGSs in total) except for Factor XII (P=1.2 ×10^−4^). The predicting *r^2^* of the 12 traits range from 0.008 to 0.105, which can also be loosely interpreted as the percentage of variance explained by the PGS. The composite scores S^WTSUM^, which were calculated from the 10 most predictive correlation-pruned PGSs, were able to improve the prediction power for all the 12 VISP traits, especially in four traits (DBP, B6, folate, and factor XII) where the prediction power was improved by more than 5-fold.

**Table 1:**
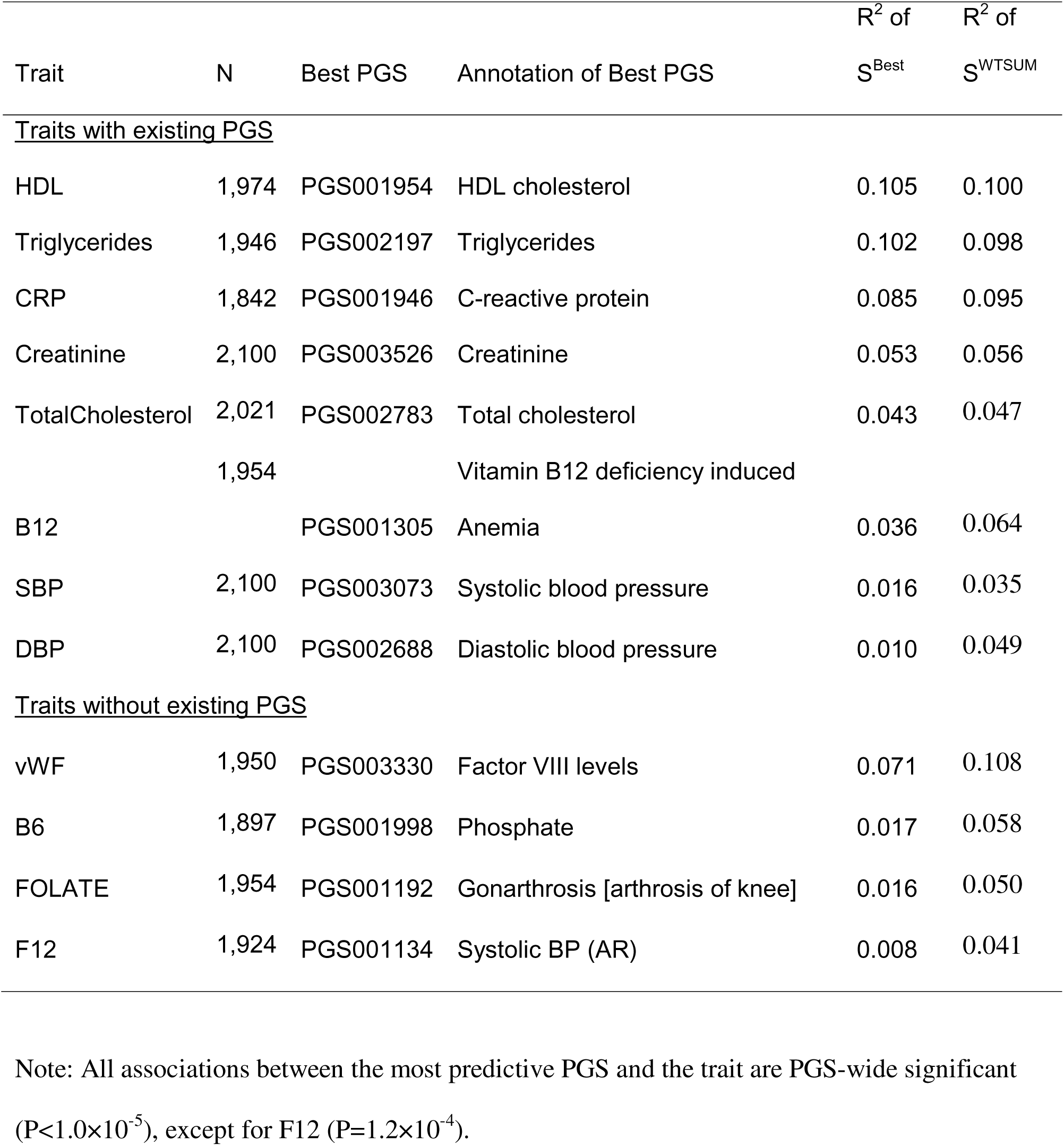
Most predictive PGSs from 12 PGS-wide scans in 2,100 VISP samples.

|  |  |  |  | R<sup>2</sup> of | R<sup>2</sup> of |
| --- | --- | --- | --- | --- | --- |
| Trait | N | Best PGS | Annotation of Best PGS | S<sup>Best</sup> | S<sup>WTSUM</sup> |
| <u>Traits with existing PGS</u> |  |  |  |  |  |
| HDL | 1,974 | PGS001954 | HDL cholesterol | 0.105 | 0.100 |
| Triglycerides | 1,946 | PGS002197 | Triglycerides | 0.102 | 0.098 |
| CRP | 1,842 | PGS001946 | C-reactive protein | 0.085 | 0.095 |
| Creatinine | 2,100 | PGS003526 | Creatinine | 0.053 | 0.056 |
| TotalCholesterol | 2,021 | PGS002783 | Total cholesterol | 0.043 | 0.047 |
|  | 1,954 |  | Vitamin B12 deficiency induced |  |  |
| B12 |  | PGS001305 | Anemia | 0.036 | 0.064 |
| SBP | 2,100 | PGS003073 | Systolic blood pressure | 0.016 | 0.035 |
| DBP | 2,100 | PGS002688 | Diastolic blood pressure | 0.010 | 0.049 |
| <u>Traits without existing PGS</u> |  |  |  |  |  |
| vWF | 1,950 | PGS003330 | Factor VIII levels | 0.071 | 0.108 |
| B6 | 1,897 | PGS001998 | Phosphate | 0.017 | 0.058 |
| FOLATE | 1,954 | PGS001192 | Gonarthrosis [arthrosis of knee] | 0.016 | 0.050 |
| F12 | 1,924 | PGS001134 | Systolic BP (AR) | 0.008 | 0.041 |

Note: All associations between the most predictive PGS and the trait are PGS-wide significant
( $P < 1.0 \times 10^{-5}$ ), except for F12 ( $P = 1.2 \times 10^{-4}$ ).

Table 2 shows the most predictive PGS results identified from each of the 12 PGS-wide scans in 377 non-European VISP samples. Majority of the samples in this subset of data were of African and Hispanic ancestry. With much fewer samples and more diverse ancestry than the full set of the VISP data, the prediction *r^2^* for each trait is not substantially reduced. However, only 5 out of the 8 traits were predicted correctly, with the same-trait-related PGSs being identified as the most predictive predictors. For the other 3 traits, including B12, SBP, and DBP, the top predictors were closely related to the underlying biology of the trait. The autoimmune disease PGS predicts B12 with *r^2^*=0.048, the stroke PGS predicts SBP with *r^2^*=0.035, and the QRS duration PGS predicts DBP with *r^2^*=0.036. The prediction improvement of the composite score S^WTSUM^ over the best PGS in samples of non-European ancestry was much higher than those of European ancestry.

**Table 2:** Most predictive PGSs from 12 PGS-wide scans in 377 non-European VISP samples.

|  |  |  |  | R<sup>2</sup> of | R<sup>2</sup> of |
| --- | --- | --- | --- | --- | --- |
| Trait | N | Best PGS | Annotation of Best PGS | S<sup>Best</sup> | S<sup>WTSUM</sup> |
| <u>Traits with existing</u> |  |  |  |  |  |
| <u>PGS</u> |  |  |  |  |  |
| HDL | 353 | PGS003534 | HDL cholesterol | 0.106 | 0.232 |
| Triglycerides | 349 | PGS003811 | Triglycerides | 0.098 | 0.226 |
| CRP | 327 | PGS000314 | C-reactive protein | 0.072 | 0.258 |
| Creatinine | 377 | PGS003246 | Creatinine | 0.036 | 0.224 |
| TotalCholesterol | 354 | PGS002783 | Total cholesterol | 0.079 | 0.221 |
| B12 | 348 | PGS002359 | Autoimmune disease | 0.048 | 0.252 |
| SBP | 377 | PGS002259 | Stroke | 0.035 | 0.187 |
| DBP | 377 | PGS002166 | QRS duration | 0.036 | 0.184 |
| <u>Traits without existing PGS</u> |  |  |  |  |  |
| vWF | 347 | PGS003330 | Factor VIII levels | 0.103 | 0.252 |
| B6 | 334 | PGS002829 | Alcohol consumption | 0.030 | 0.197 |
| FOLATE | 349 | PGS004340 | SHBG | 0.041 | 0.238 |
| F12 | 335 | PGS002633 | Cardiovascular disease | 0.047 | 0.248 |

Figure 1 shows the prediction performance of the 5 PGS models developed by 5 PGS methods including C&T, Lassosum2, LDpred2, SECRET-Best, and SECRET-WTSUM. Across all 12 traits, the 2 SECRET methods substantially outperform the 3 existing methods. *E.g.*, for traits with a PGS that explain ∼10% of the phenotypic variation (according to the SECRET *r^2^* estimates, which includes HDL, Triglycerides, and CRP), none of the existing methods were able to predict with *r^2^*>0.02. In this cross-validation prediction analysis, several of the SECRET results in Table 1 (*i.e.*, without validation) were validated here. Note the *r^2^* of S^Best^ in the training set (*i.e.*, without validation) was indicated in dashed line. Compared to the *r^2^* of S^Best^ in the validation set, the two *r^2^*’s were comparable without substantial bias (the latter is larger in 5 out 12 times). The observation that SECRET-WTSUM outperformed SECRET-Best in Table 1 were validated in 3 out of 4 traits (B6, FOLATE, and F12). In addition, SECRET-WTSUM outperformed SECRET-Best for traits B12 and SBP.

**Figure 1:**
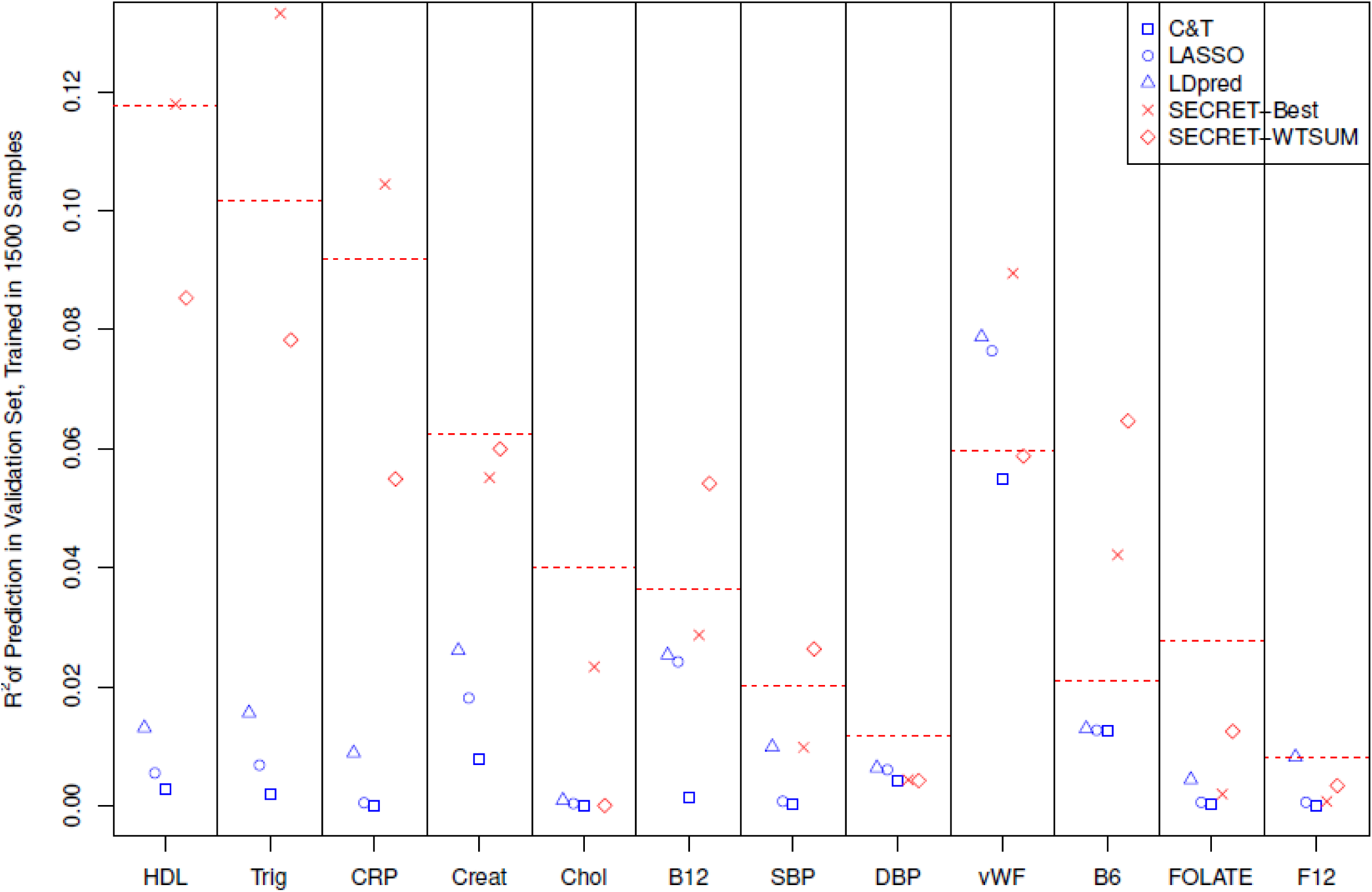
Comparison of prediction performance of 5 methods for developing PGS models. For each PGS method, a PGS model was developed for each of the 12 traits using a training set consisting of 1,500 samples, and the prediction *r^2^* of this PGS model was calculated in an independent validation set consisting of 223 samples. The prediction *r^2^* for the 3 existing PGS methods and the 2 SECRT methods are indicated in blue and red respectively in the figure. Each of the horizontal dashed lines in red is the prediction *r^2^* of a SECRET-Best PGS that was calculated in the training set without any validations.

### Data application: Family-based PGS scan in T1DGC

We incorporated a family-based PGS method into SECRET and benchmarked it utilizing the T1DGC family-dataset^28^. In the T1DGC PGS-wide scan, we found that among the top 30 PGS predictors, 23 PGSs were developed for T1D, and the other 7 were T1D-related (e.g., “age diabetes diagnosed”). The best individual PGS that predicts T1D was PGS004162 (T1D)^29^, with predicting AUC=0.776, which is somewhat lower than a population-based T1D prediction^1,2^. Figure 2 lists the top 30 PGS predictors of T1D in our PGS-wide scan.

**Figure 2:**
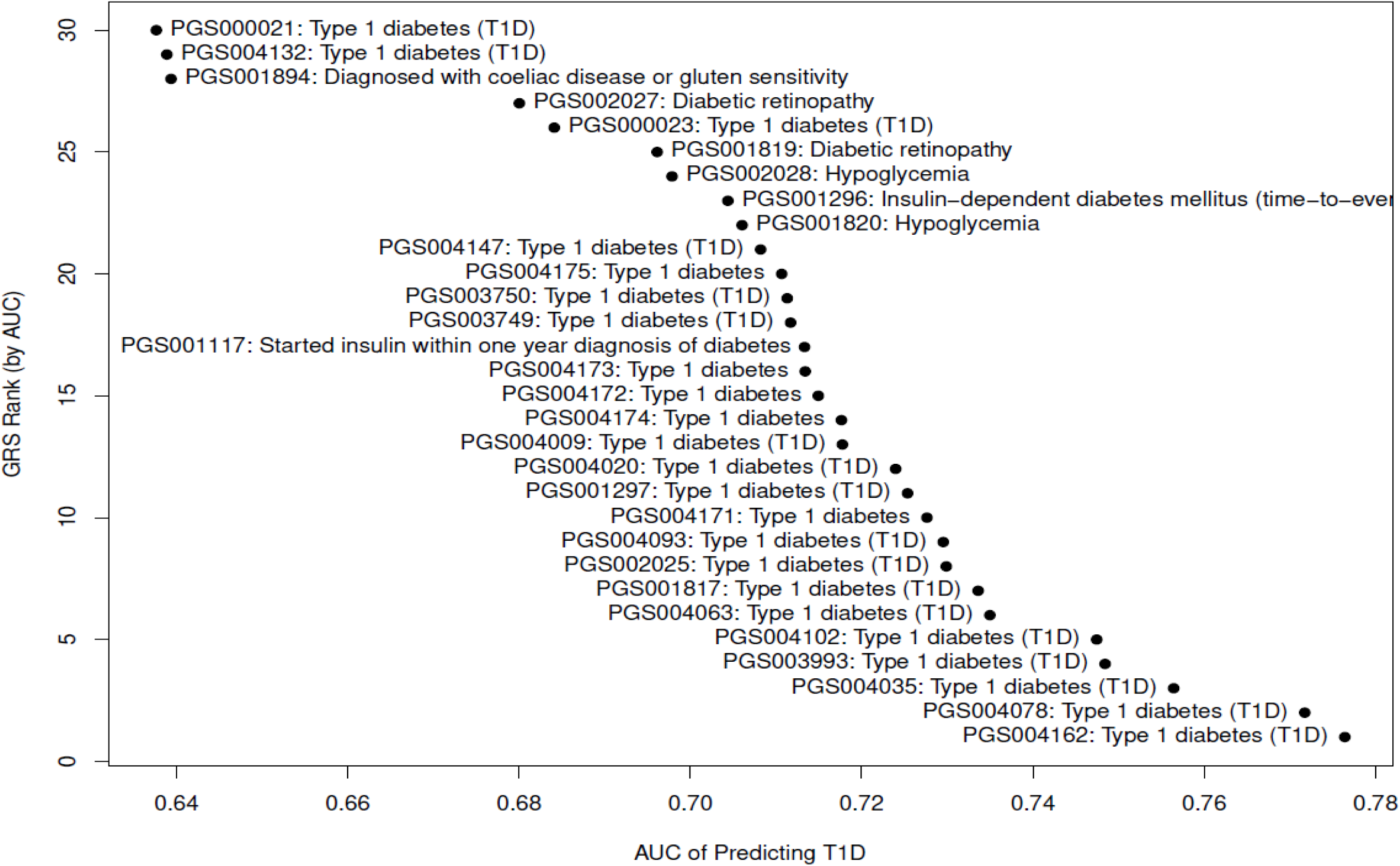
Top 30 PGS predictors of T1D in T1DGC families. The ranks of the PGS predictors are based on the AUC of prediction, and the best predictors are located at the bottom.

The MHC region located on chromosome 6 harbors half of the genetic risk for T1D, and therefore we screened MHC and non-MHC PGSs. The best PGS predictors of T1D using MHC PGSs or non-MHC PGSs were two distinct T1D PGSs (PGS001817 and PGS003993), each with AUC=0.72. Other than the T1D PGSs, T2D (type 2 diabetes) PGSs also predicted T1D, but only when the PGS was constructed using SNPs inside the MHC region. Figure 3 shows the role of MHC region in using T2D PGSs to predict T1D. Across both males and females, the T2D PGSs that were constructed using SNPs in the MHC region could predict T1D with AUC as large as 0.70 in males and 0.68 in females. In contrast, none of the T2D PGSs outside of the MHC region could predict T1D (AUC < 0.53).

**Figure 3:**
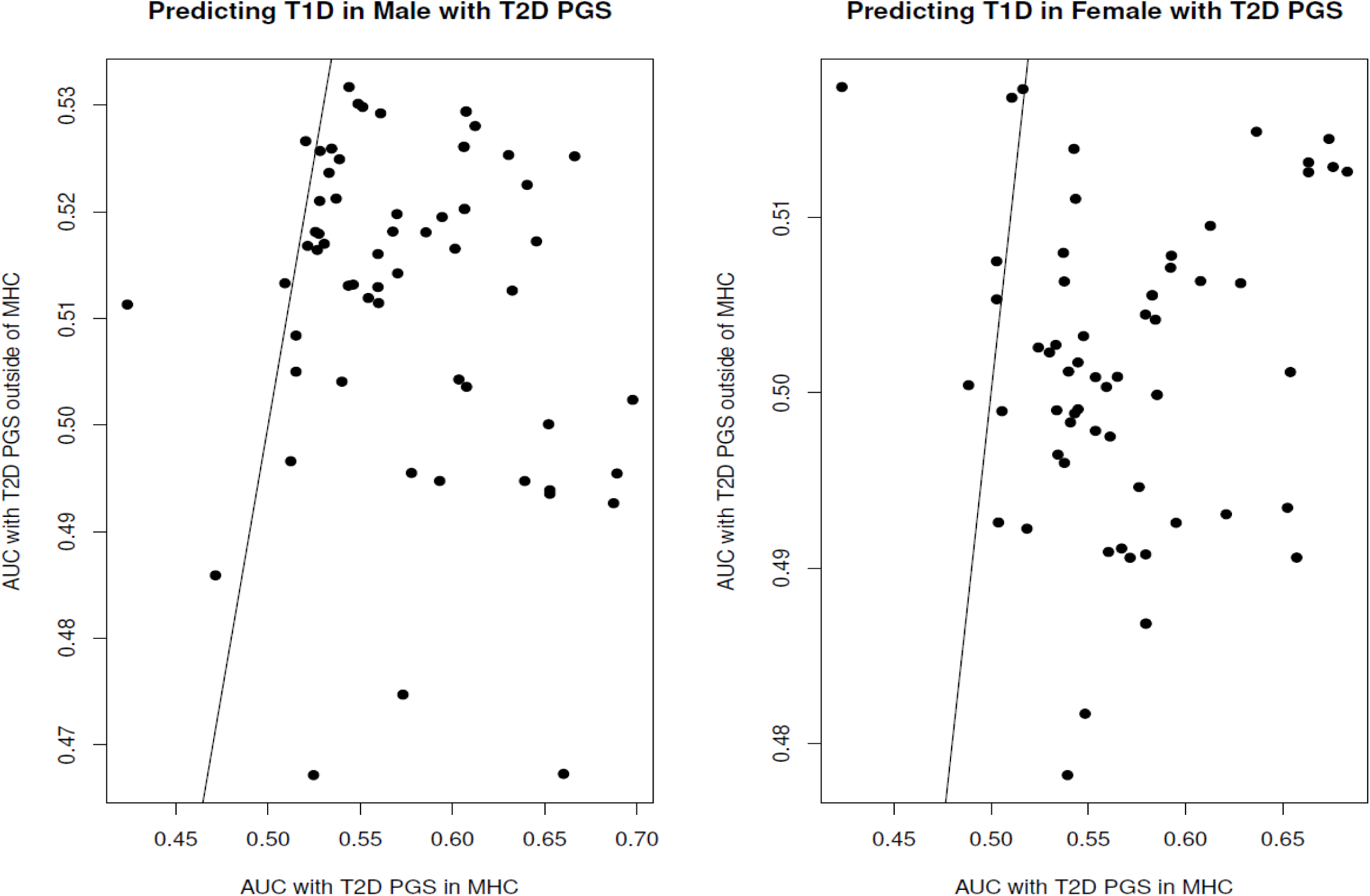
Potential role of MHC region in using T2D PGS to predict T1D risk. Each T2D PGS was partitioned into an MHC-PGS that was calculated using the MHC SNPs only, and a non-MHC-PGS that was calculated using the SNPs outside of the MHC region. The AUC was calculated separately for each of the portioned PGS, for all T2D PGS models, and the AUC for the MHC-PGS was plotted against an AUC for the non-MHC-PGS for each of the T2D PGS models, in males and in females only.

### Sample sizes and powers in simulations

Simulations using the VISP and T1DGC datasets indicated hundreds of samples may be sufficient to identify a correct PGS model for a trait using the SECRET method (Figure 4). For traits such as T1D, a sample size as small as 100 could be sufficient, with a power of 89% to identify a T1D PGS as the best PGS. For traits with a PGS that explain ∼10% of phenotypic variation, such as HDL, Triglycerides, and CRP, 400 samples would have >80% power to correctly identify the trait-related PGS as the best predictor in SECRET.

**Figure 4:**
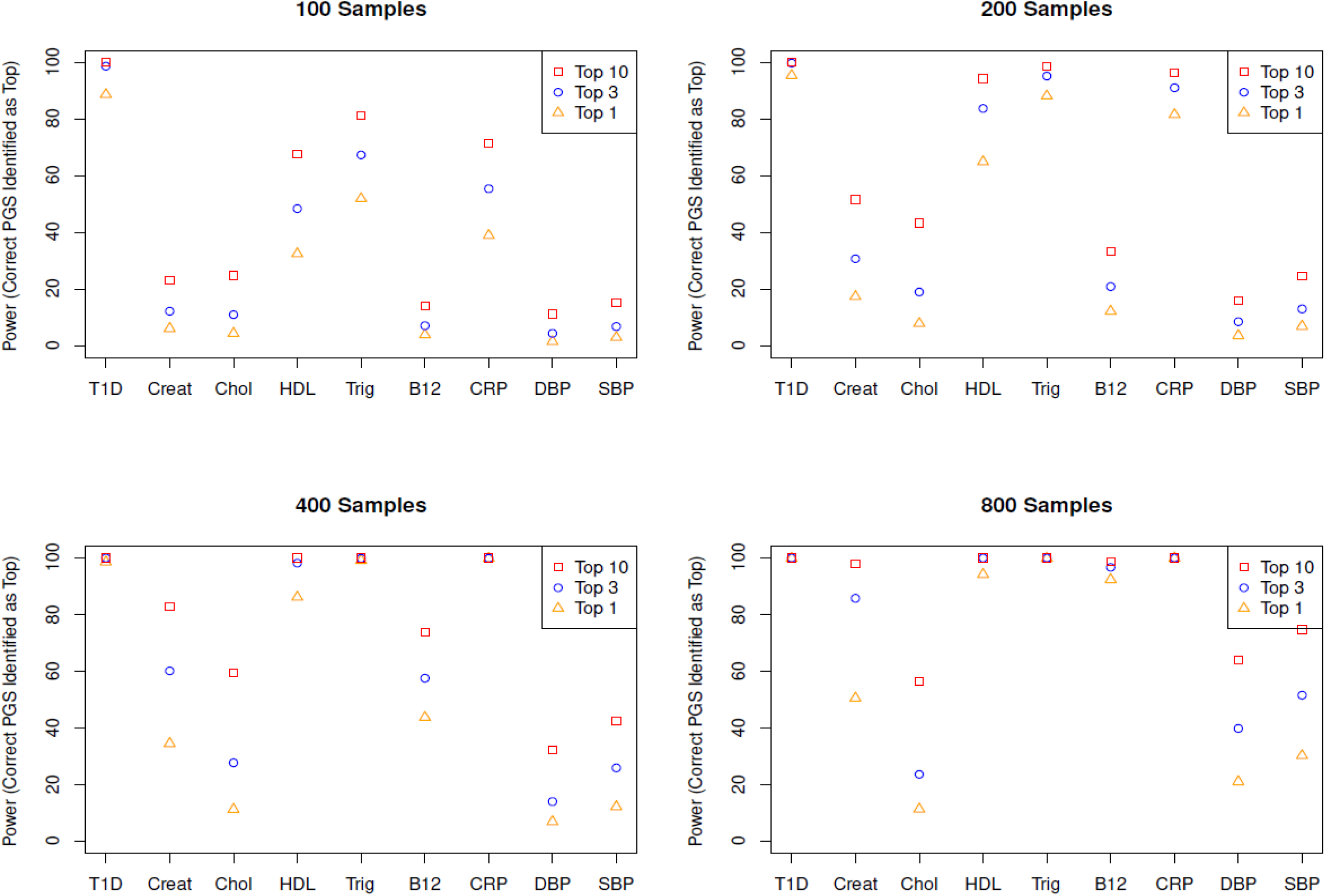
Power of identifying the correct PGS using the SECRET-Best method for 9 traits in a simulation study. In each round of simulation, we randomly selected a subset of samples (N=100, 200, 400, and 800 respectively) for 8 traits from VISP and T1D from the T1DGC family data. Note for the family data, the random selection was by families instead of by individuals. We carried out a PGS-wide scan using the SECRET-Best method and evaluated if a PGS model for the trait of interest could be identified as one of the top-ranked PGS models. We repeated this procedure 1,000 times and calculated the power of identifying the correct PGS for the trait.

## Discussion

We present a PGS method that leverages public resources such as the PGS Catalog for developing predictive PGS models with as few as hundreds of samples, much lower than any existing methods. This method can be especially valuable to genetic studies of not well-studied traits such as treatment and drug responses.

Using a modest-sized dataset that is not part of the PGS Catalog, we demonstrate that SECRET can effectively screen and identify the most relevant PGS for traits that already have an entry in the PGS Catalog. As the PGS catalog continues to expand rapidly, we expect more traits will have their own specific PGS model to be the best predictor. For traits that do not have a PGS model in the PGS Catalog currently, SECRET can identify a genetically closely-related PGS model, where the corresponding trait serves as a risk factor, or shares a common biological pathway. In addition, the two PGS models developed by the SECRET methods may have substantially higher prediction power than any of the existing PGS methods.

Like a GWAS scan, the PGS-wide scan proposed as part of the SECRET approach screens and identifies PGS models with the strongest association with the trait. Although the proposed SECRET framework does not incorporate inference of causality, the PGSs for true risk factors would not be missed in a PGS-wide scan when they are present in the PGS Catalog (assuming the population structure and other covariates have been properly adjusted). As a result, a major feature of SECRET is its utility in discovery or prioritization of trait-related risk factors. For clinical applications where causal inference is important (e.g., drug/treatment response studies, or clinical trials), the top PGS predictors identified through SECRET can be validated, statistically, through Mendelian Randomization (MR) analysis using the GWAS data.

A family design has the advantage of eliminating confounding factors such as population stratification and assortative mating, which present a great challenge to many PGS applications^30^. Our T1D PGS scan successfully identified T1D PGSs as the top predictors and systematically evaluated the predictive power of dozens of T1D PGSs, highlighting the potential of family data to enhance effectiveness of PGS applications.

We demonstrated that a large number of T2D PGS models using only MHC variants can predict T1D status (with AUC as large as 0.70). However, it is still not clear to us whether it indicates T1D and T2D may share the same causal pathway, or it indicates potential data contamination during the PGS model development. Note the T2D models that predict T1D are all from large biobanks, and a small portion of T1D patients that are mis-reported as T2D could lead to this prediction result. Further analysis beyond SECRET is needed for clarification.

In addition to identifying the most predictive PGS models, we also implement a method that further improves the prediction power by combining multiple PGS models. Although this method is fast and built into SECRET, there may exist alternative methods^31^ that outperform our method. We have not systematically compared various methods to combine multiple PGS models, which is a future direction of our work. An optimal strategy can be to first generate the top PGS models and scores using SECRET, and then to combine the top PGSs using the best available method.

In summary, the method and tool developed in SECRET provide a valuable resource for researchers to generate predictive PGSs in studies with limited sample sizes, enable systematic evaluation of PGSs, and stimulate insights and hypotheses about their genetic and biological underpinnings.

## Electronic Database Information

https://www.kingrelatedness.com/SECRET/ for the implementation of algorithm for SECRET

